# Getting more from heterogeneous HIV-1 surveillance data in a high immigration country: estimation of incidence and undiagnosed population size using multiple biomarkers

**DOI:** 10.1101/345710

**Authors:** Federica Giardina, Ethan Romero-Severson, Maria Axelsson, Veronica Svedhem, Thomas Leitner, Tom Britton, Jan Albert

## Abstract

**Background:** Most HIV infections originate from individuals who are undiagnosed and unaware of their infection. Estimation of this quantity from surveillance data is hard because there is incomplete knowledge about *i)* the time between infection and diagnosis (TI) for the general population and *ii)* the time between immigration and diagnosis for foreign-born persons.

**Development:** We developed a new statistical method for estimating the number of undiagnosed people living with HIV (PLHIV) and the incidence of HIV-1 based on dynamic modeling of heterogenous HIV-1 surveillance data. We formulated a Bayesian non-linear mixed effects model using multiple biomarkers to estimate TI accounting for biomarker correlation and individual heterogeneities. We explicitly model the probability that an HIV-1 infected foreign-born person was infected either before or after immigration to distinguish between endogenous and exogeneous incidence. The incidence estimator allows for direct calculation of the number of undiagnosed persons.

**Application:** The model was applied to surveillance data in Sweden. The dynamic biomarker model was trained on longitudinal data from 31 treatment-naïve patients with well-defined TI, using CD4 counts, BED serology, polymorphisms in HIV-1 *pol* sequences, and testing history. The multiple-biomarker model was more accurate than single biomarkers (mean absolute error 1.01 vs ≥ 1.95). We estimate that 813 (95% CI 780-862) PLHIV were undiagnosed in 2015, representing a proportion of 10.8% (95% CI 10.4-11.3%) of all PLHIV.

**Conclusions:** The proposed methodology will enhance the utility of standard surveillance data streams and will be useful to monitor progress towards and compliance with the 90-90-90 UNAIDS target.

**Key messages:**

- Combined heterogeneous HIV-1 surveillance data and biomarker data can be used to estimate both local incidence and the number of undiagnosed people living with HIV.
- Explicit modeling of the dynamics, heterogeneity, and correlation of multiple biomarkers over time improved estimation of time between infection and diagnosis.
- Explicit modeling of the probability that foreign-born persons were infected before or after immigration improves accuracy of estimates of endogenous incidence and undiagnosed persons living with HIV.
- The endogenous incidence of HIV-1 in Sweden is declining, despite continued immigration of HIV-1 infected persons.
- The proportion of undiagnosed PLHIV decreased over 2010-2015 and was estimated to be 10.8% (95% CI, 10.4-11.3%) in 2015.

## Introduction

The majority of new HIV-1 infections originate from individuals who are undiagnosed and unaware of their infection, especially in countries with good access and adherence to antiretroviral therapy (ART) ^1–4^. Thus, knowledge about the size of the undiagnosed HIV-1 population is highly relevant for public health and HIV prevention. In 2014, the Joint United Nations Programme on HIV and AIDS (UNAIDS) launched the 90-90-90 target, which states that by 2020 i) 90% of all persons living with HIV (PLHIV) should know their status ii) 90% of all diagnosed HIV cases should receive ART and iii) 90% of all people receiving ART should have achieved viral suppression ^5^. The European Center for Disease Prevention and Control (ECDC) estimates that 1 in 7 (14%) PLHIV in Europe are unaware of their infection ^6^.

Estimation of the size of the undiagnosed population is challenging because the time of infection usually is unknown and the time until diagnosis (TI) is highly variable due to differences in the testing behavior, risk awareness, and rate of disease progression. Until now, most estimates of HIV-1 incidence and undiagnosed population have been based on methods that classify patients as recently or long-term infected rather than estimate TI directly. Several such methods based on CD4+ T-lymphocyte (CD4) counts, HIV-1 antibody tests and viral sequence diversity have been described. CD4 counts are commonly used ^7^, but their rate of decline is variable ^8–10^ limiting their utility as a single biomarker. HIV-1 antibody concentrations increase over time from infection approaching an asymptote in long-term infections ^11^. The BED IgG-capture enzyme immunoassay (BED assay) has been used to estimate incidence ^12,13^ by classifying recent and long-term infections. Sequence-based methods exploit the increase in intra-patient sequence diversity following infection, which can be approximated by the fraction of polymorphic nucleotides in the partial HIV-1 *pol* gene sequences that are used for detecting drug resistance mutations ^14,15^.

Fewer studies have attempted to estimate TI continuously, rather than distinguishing between recent and long-term infections, and to account for individual variations. Sommen et al. ^16^ used antibody levels to two HIV-1 antigens (IDE and V3) to calculate the posterior distribution of HIV-1 infection times and incidence in France. Romero-Severson et al. ^11^ used BED assay results to combine a model of within-host time-continuous IgG dynamics ^17^ with a Bayesian estimator for the incidence of HIV-1 in Sweden in 2002-2009. However, these studies neither explicitly accounted for immigration of infected persons and endogenous infection of immigrated persons, nor attempted to estimate the size of the undiagnosed HIV-1-infected population.

Here we present a statistical method to estimate HIV-1 incidence and the number of undiagnosed people living with HIV (PLHIV). The method is based on *i)* a model of the joint dynamics of multiple-biomarkers from the time of infection that allow a more accurate estimation of individual TIs and *ii)* the estimation of the time from immigration to diagnosis for exogenous infections using unlinked surveillance data. The TI model uses BED, CD4 and *pol* polymorphisms, but can be easily generalized to include other or additional biomarkers. The estimation of the time from immigration to diagnosis allows the assessment of how endogenous and exogeneous infections contribute to the undiagnosed fraction.

In the application of the model to surveillance data from Sweden we estimated that endogenous HIV-1 infections have decreased over 2010-2015 and that 10.8% (95% CI, 10.4-11.3%) of all infected persons in Sweden were undiagnosed in 2015. This estimate is in line previously published results^18^ and allowed uncertainty quantification.

## Methods

### Multiple-biomarker model for estimation of time of infection

The *K* biomarkers 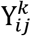, *k* = 1, …,*K* are modelled jointly as a function of TI as follows:

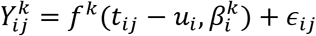

where *i* denotes the individual with biomarkers measured at calendar time *t*_*ij*_ after infection date *u*_*i*_. The dynamics are described by biomarker specific curves *f* ^*k*^(·), fixed and random effects 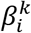 and 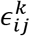 as *i.i.d*. error terms such that 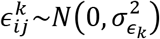. The model is formulated in a Bayesian framework and fitted to longitudinal data with known TI using Markov chain Monte Carlo (MCMC). For details on the prior distributions assigned to the unknown parameters, see Supplementary Data, Section 1.1.

The Bayesian formulation allows us to use this model to estimate directly the unknown time of infection for new individuals, by treating individual TI as “latent” variables and assigning them a prior distribution. This approach was used in a leave-one out cross validation analysis using the same longitudinal dataset (where TI was represented by any of the *t*_*ij*_ − *u*_*i*_), as well as in the prediction of TI for newly infected individuals, with biomarker measured only at time of diagnosis, i.e. *j* = 1.

### Handling of foreign-born cases

To estimate HIV-1 incidence and undiagnosed PLHIV, foreign-born persons infected before first arrival should only be counted after arrival. Immigration dates are rarely reported in HIV surveillance systems alongside HIV biomarkers information. For that, we first estimate a typical distribution for the time interval between immigration *e* and HIV diagnosis using independent and unlinked surveillance data.

### Incidence estimation

We extended the model proposed by Sommen et al. ^16^ to estimate HIV-1 incidence. Our modifications allow us to *i)* consider all reported cases between *t* and the present time in the yearly incidence estimation, rather than only cases reported in a specific year, and *ii)* explicitly account for imported infections. We denote with *I*_[τ_1_,τ_2_]_ the incidence in time period [τ_1_, τ_2_], which will typically be one year, and with τ_3_ the end of the observation period, e.g. present time. The incidence is expressed as a weighted sum of posterior densities of infection (or entry) times:

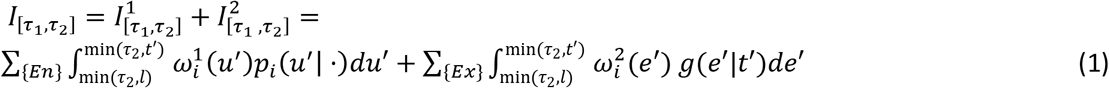

where *En* and *Ex* denote endogenous and exogenous infections diagnosed in [τ_1_, τ_3_], respectively. Here, *P*_*i*_(*u*′| ·) represents the posterior distribution of infection times for individuals diagnosed at *t*′ obtained using observed biomarkers and *g*(*e*′|*t*′) is the “backward” distribution used to generate entry times *e* for foreign-born cases given diagnosis dates *t*′ in the country (by year of diagnosis). The time of the latest negative test that individual *i* may have is denoted by *l*. The second term in (1) is evaluated only when the generated immigration time is more recent than the infection time as estimated by the biomarkers. Full definitions of the weights 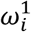 and 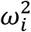 can be found in the Supplementary Data.

### Estimation of undiagnosed fraction

The calculation of the HIV-infected undiagnosed individuals in a specific year follows directly as the product of the estimated HIV-incidence by the probability of not being diagnosed by the end of the same year, stratified by transmission route and country of infection (endogenous or exogenous infection). In particular, to calculate the number of undiagnosed individuals *U*_τ_3__ at time τ_3_ we use: 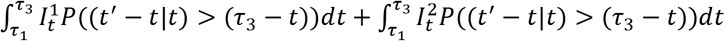 where 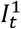 and 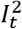 are the two terms defined in (1) and *t*′ are the diagnosis times for individuals estimated to have been infected (or immigrated) at time *t*. Credible intervals are obtained by generating 100 bootstrap samples of size 1000 from the distributions of infection times. All statistical analyses were carried out using R version 3.3. ^19^ and JAGS ^20^. For further details on the model and parameters see the Supplementary Data, Section 1 and Section 3.

### Surveillance data

The methods were applied to estimate the incidence of infection and the number of undiagnosed in Sweden. Surveillance data in Sweden include multiple data sources. We used published data on 1,357 HIV-1 infected patients diagnosed in Sweden between 2003 and 2010 ^11,17,21,22^ with complete data on CD4 concentration, BED level, and *pol* polymorphism counts at diagnosis. These patients represented 39% of all patients diagnosed in Sweden during this period. These data included likely country of infection, transmission route, last negative and first positive HIV-1 test, laboratory evidence of primary HIV-1 infection, plasma HIV-1 RNA levels, but not data on time of arrival in Sweden for foreign-born persons.

The biomarker model used data from three biomarkers (BED, CD4 and *pol* polymorphisms) and was trained on a subset of 31 treatment-naïve patients having longitudinal biomarker data and known infection times. For comparison, the three biomarkers were also modelled separately. To assess the representativeness of the dataset, we compared the estimated parameter values to values reported in the literature.

Data on the yearly number of diagnosed HIV-1 cases in Sweden between 2003 and 2015, stratified by reported transmission route, country of infection (Sweden or abroad), and AIDS at HIV diagnosis are collected as part of mandatory national case reporting. In total 5,777 patients were diagnosed in Sweden during 2003-2015. For 2,466 of 2,978 (83%) of the foreign-born patients, data on the time between first arrival and diagnosis in Sweden was available; however, these data were anonymized and could not be linked to the other data.

For details on the patients and laboratory methods, see Table S1 and Section 2 of the Supplementary Data.

## Results

### The multiple-biomarker model improved estimation of TI

The parameter values describing the growth or decline of the biomarkers for the training data (Supplementary Data, Figure S1-S3 and Table S2) agreed with published values ^7,14,23–25^, which justified the use of the model on biomarker data from other HIV-1 patients. A leave-one-out cross-validation analysis showed that the multiple biomarker model gave more accurate estimates of TI than each of the single biomarkers according to four different measures of precision (Table 1). TI estimates were more precise if biomarkers were measured shorter after infection (MAE=0.91 at <1 year; 1.20 at 1-2 years; 1.40 at >2 years, see Table S3 in the Supplementary Data). Biomarker measurements at two or three time points, rather than a single time point, only slightly improved TI estimation (Table S4 in the Supplementary Data). This motivated the use of single time point biomarker measurements in the application to newly diagnosed patients in Sweden.

**Table 1.** Mean predictive performance of single and multiple biomarker models assessed with a leave-one-out cross validation analysis evaluated by four measures of precision. CD4, CD4+ T-lymphocyte count; BED, antibody levels measured using the BED IgG-Capture enzyme immunoassay model; POL, fraction of polymorphisms in HIV-1 *pol* gene sequences; MBM, multiple biomarker model based on CD4, BED and *pol*; MAE, mean absolute error; RMSE, root mean square error.

| Model | Bias | MAE | RMSE | Coverage |
| --- | --- | --- | --- | --- |
| CD4 | -2.62 | 3.12 | 3.77 | 0.81 |
| BED | -1.80 | 1.95 | 2.21 | 0.90 |
| POL | -2.16 | 2.22 | 3.20 | 0.81 |
| MBM | -0.68 | 1.01 | 1.38 | 0.93 |

### Time between infection and diagnosis varied by transmission route

Figure 1 shows the distribution of the estimated time until diagnosis in the three main transmission groups (MSM, IDU and HET) broken down by country of origin (Sweden or abroad). For MSM and IDU the estimated time between infection and diagnosis showed similar distributions, with around 60% of individuals being diagnosed within 1 year after infection and 8% being diagnosed >5 years after infection. Heterosexually infected persons had longer time until diagnosis with around 31% and 19% being diagnosed within 1 year and after >5 years after estimated TI, respectively. As expected, persons reported to have been born abroad also had longer time between estimated TI and diagnosis in each transmission group, as some of them are likely to be exogenous infections (overall median 2.0 years vs 0.89 years, Mann-Whitney test, P<0.05).

**Figure 1.**
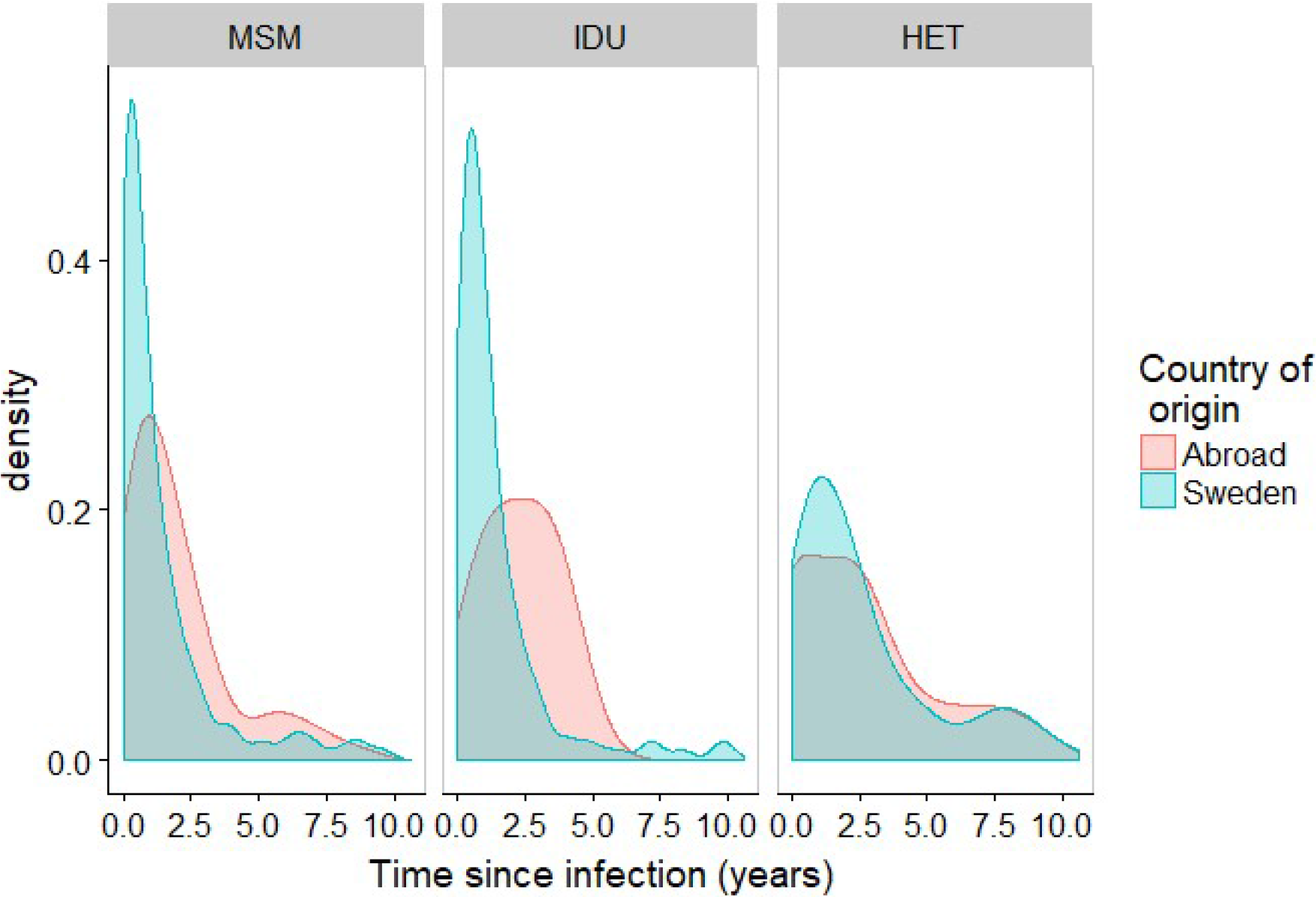
Distribution of estimated time from infection (TI) to diagnosis by transmission route and country of origin modeled using CD4, BED and *pol* polymorphism data obtained at or close to diagnosis from 1,357 patients diagnosed in Sweden 2003-2010. MSM, men who have sex with men; IDU, intravenous drug users; HET, heterosexual transmission route.

### Decreasing HIV-1 incidence in Sweden

To estimate HIV-1 incidence in Sweden we explicitly modeled the impact of migration so that patients estimated to have been infected before immigrating to Sweden contributed to incidence only from the estimated date of first entry into the country (Figure 2). Figure 3 shows that the incidence of endogenous infections decreased in all three transmission groups, especially among MSM and IDU. We estimate that 123 (95% CI, 105-139) infections occurred in Sweden in 2015 (31 MSM, 3 IDU and 89 HET), compared to 280 (95% CI, 258-306) in 2010 (109 MSM, 12 IDU and 159 HET), P<0.01. In contrast, the incidence of persons entering Sweden already infected (i.e. exogenous infections) was estimated to have been stable or slightly increasing; with 257 (95% CI, 200-307) persons in this category in 2015 (78 MSM, 13 IDU and 166 HET), compared with 242 (95% CI, 221-262) in 2010 (P=0.15).

**Figure 2.**
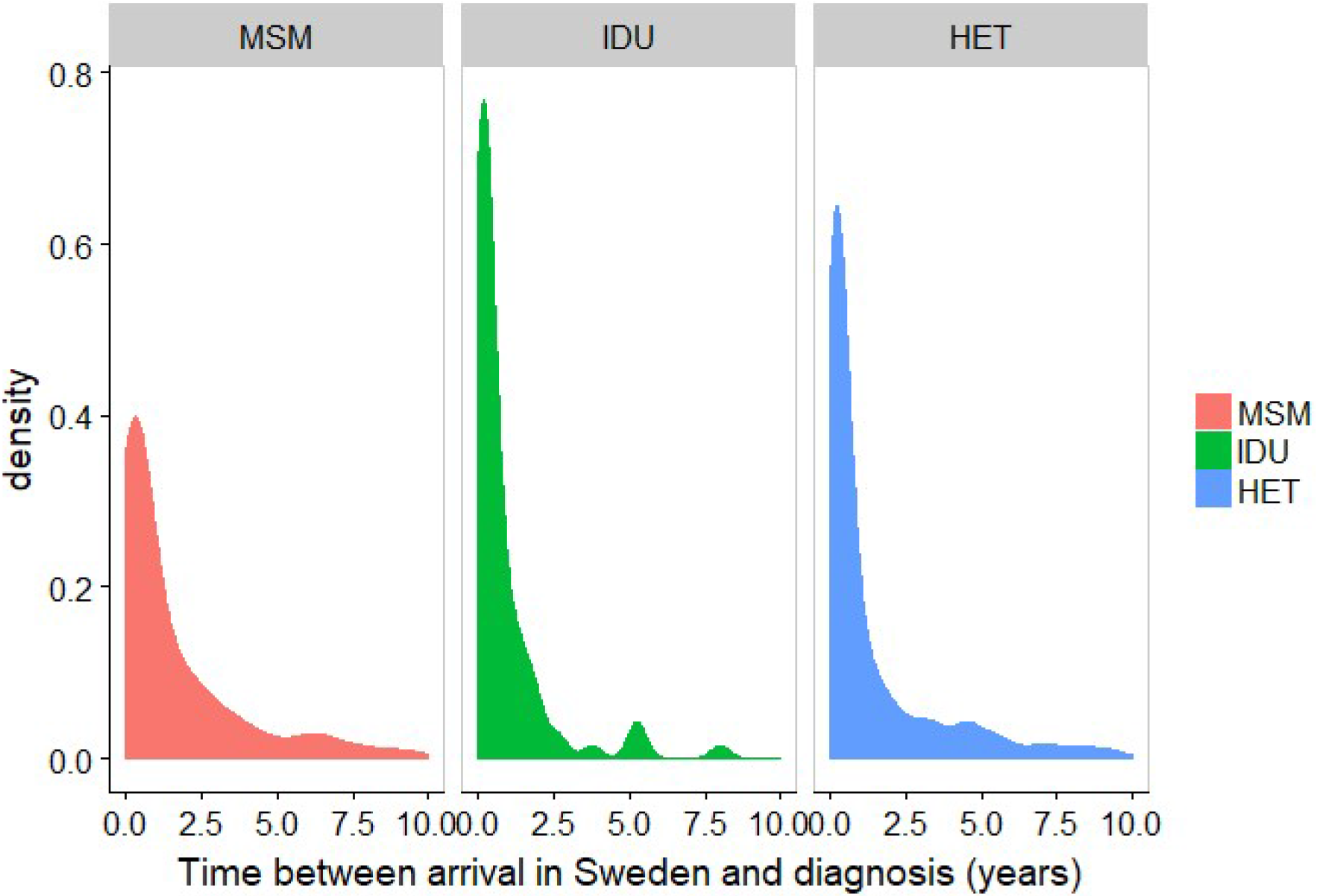
Distribution of estimated time from arrival in Sweden to diagnosis by transmission route modeled using data on time between first entry into Sweden and diagnosis for 2,466 patients obtained from the Public Health Agency of Sweden. MSM, men who have sex with men; IDU, intravenous drug users; HET, heterosexual transmission route.

**Figure 3.**
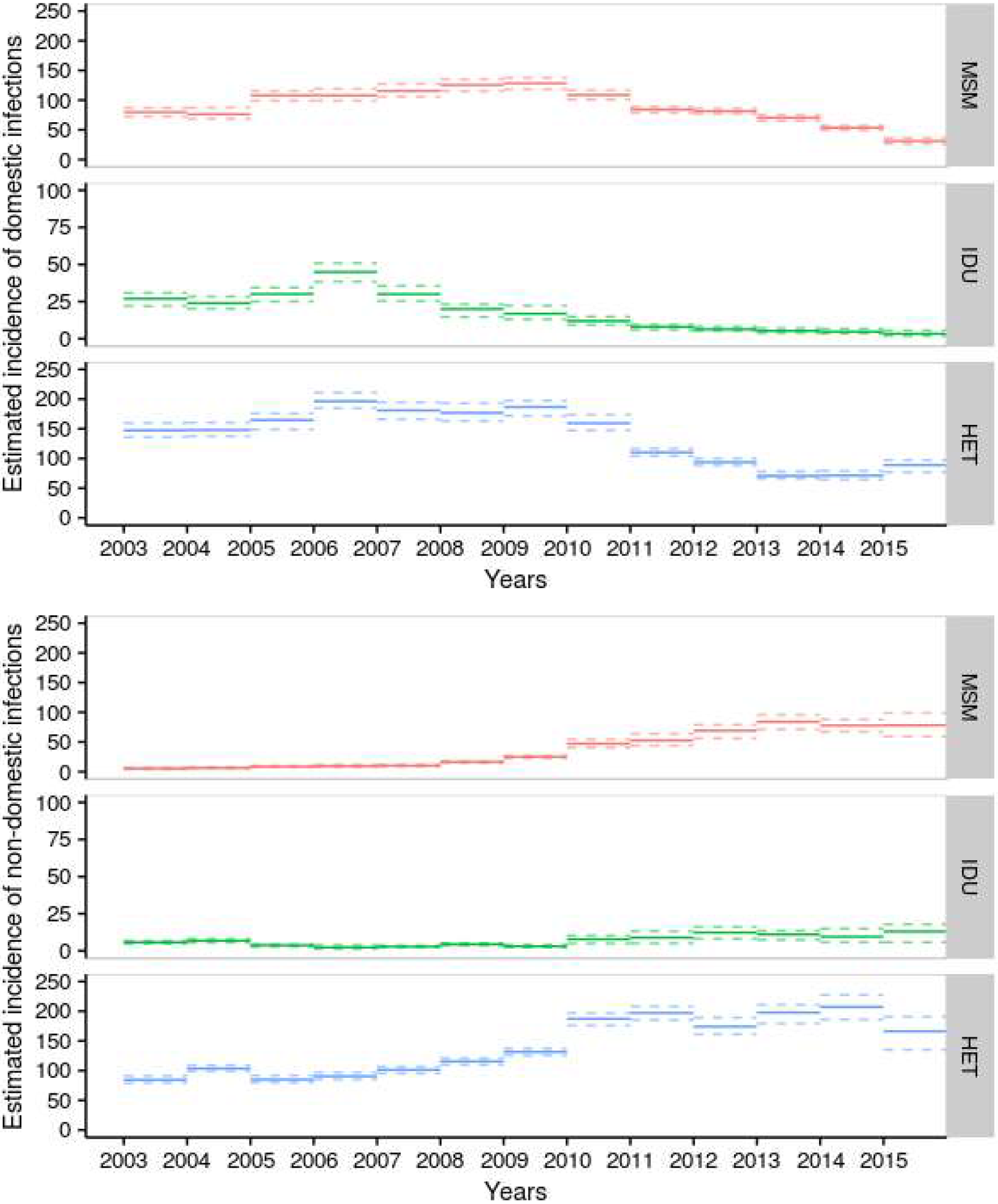
Estimated HIV-1 incidence in Sweden 2003-2015 per year and transmission route. The model explicitly accounts for exogenous infections. Thus, persons estimated to have been infected before first entry in Sweden only contribute to incidence in Sweden from the estimated date of entry. The upper panels show estimated incidence of HIV-1-infection among persons residing in Sweden. The lower panels show the incidence of HIV-1-infected persons entering Sweden for the first time. MSM, men who have sex with men; IDU, intravenous drug users; HET, heterosexual transmission route. Note the scale on the y-axis is different for IDU.

### Proportion of undiagnosed HIV-1-infection persons close to 10% UNAIDS target

Our model also provided estimates on the number of undiagnosed PLHIV in Sweden at the end of the years 2010 to 2015. Table 2 shows that number of undiagnosed PLHIV in Sweden was estimated to have decreased from 907 (95% CI, 875-940) persons in 2010 to 813 (95% CI, 780-862) in 2015. During the same period the number of diagnosed patients linked to care increased from 5281 to 6747, which means that the proportion of undiagnosed PLHIV decreased from 14.7% (95% CI, 14.2-15.1%) in 2010 to 10.8% (95% CI, 10.4-11.3%) in 2015. Of the undiagnosed PLHIV in 2015, 340 persons were estimated to have been infected while living in Sweden (78 MSM, 8 IDU and 254 HET) and 473 persons were estimated to have been infected before first entering Sweden (164 MSM, 11 IDU and 298 HET).

**Table 2.** Estimated number and proportion of undiagnosed HIV-1-infected persons in Sweden 2010-2015. The number of undiagnosed in 2015 was calculated using the estimated incidence divided by main transmission route over a period of 13 years (2003-2015). The number of undiagnosed in previous years (2014-2010) was calculated assuming the incidence in the years before 2003 constant and equal to that of 2003. Also, the time distribution between infection and diagnosis was assumed to be transmission route dependent but year independent. PLHIV, persons living with HIV.

| Year | No. of PLHIV linked to care | No. of undiagnosed PLHIV (95% CI) | Proportion undiagnosed PLHIV (95% CI) |
| --- | --- | --- | --- |
| 2010 | 5281 | 907 (875-940) | 14.7% (14.2-15.1%) |
| 2011 | 5616 | 894 (863-927) | 13.8% (13.3-14.1%) |
| 2012 | 5918 | 877 (845-902) | 12.9% (12.5-13.2%) |
| 2013 | 6205 | 865 (834-890) | 12.2% (11.8-12.5%) |
| 2014 | 6469 | 848 (816-882) | 11.5% (11.2-11.9%) |
| 2015 | 6747 | 813 (780-862) | 10.8% (10.4-11.3%) |

## Discussion

In this manuscript we present a new method for estimating HIV incidence and size of undiagnosed population that differentiates between exogenous and endogenous infection of foreign-born cases. Our method builds on *i)* a Bayesian model to estimate time of infection (TI) using multiple biomarkers and *ii)* an extension of the incidence estimator proposed by Sommen at al.^16^ to explicitly model the probability that an HIV-1 infection in an immigrating person occurred before or after immigration. Our methodology was designed for use on heterogeneous and unlinked surveillance data.

We applied the method to data from Sweden using three biomarkers (CD4 counts, BED assay results and proportion of polymorphic sites in HIV-1 *pol* sequences as HIV-1 biomarkers) as well as data on main transmission route, presence of a last negative test, AIDS at diagnosis, diagnosed primary HIV-1 infection (PHI) to estimate TI. We treated TI as a continuous random variable rather than as a simple binary state of being recently or long-term infected, which increases power and reduces bias. We found that a combination of the three biomarkers gave more precise TI estimates than one or two of these biomarkers. Each of the three biomarkers have certain advantages and limitations and show considerable inter-individual variability. A well-known disadvantage of the BED assay is that it can give false-positive recent results for patients with low CD4 counts ^26,27^. This problem is reduced by our model because the inclusion of CD4 counts and *pol* polymorphisms partially corrects false-recent BED results, and because persons with AIDS at diagnosis where assigned a prior distribution with an average time from infection to diagnosis of 8 years ^28^ (Supplementary Data). Furthermore, low BED results in AIDS patients were rare in our dataset (Supplementary Data, Figure S4, S7 and Table S5).

We observed a decrease in the estimated incidence of HIV-1 infections and number of undiagnosed PLHIV in Sweden. The proportion of undiagnosed PLHIV was estimated to be 10.8% in 2015, which is close to the 10% UNAIDS 90-90-90 target. We estimated that the incidence of HIV-1-infections among persons residing in Sweden decreased by almost two-thirds from 2010 to 2015. The decrease was more pronounced in MSM and IDU than among heterosexually infected persons. This agrees with the fact that 87% of persons with diagnosed HIV-1-infection were on effective ART in 2015 ^18^, which means that they were effectively non-infectious ^29^, and that the time between infection and diagnosis was shorter among MSM and IDUs than among heterosexuals (Figure 2). This important distinction between endogenous and exogenous infections would not be noted if we had not accounted for immigration. Thus, the overall incidence was almost constant over the observation period (Fig S9 in Supplementary Data).

The training dataset was relatively small, consisting of longitudinal samples from 31 untreated HIV-1-infected patients. This is mainly because modern guidelines recommend treatment for all patients irrespective of immune status and the Swedish HIV-1 epidemic is relatively small. Thus, a limited number of consenting patients fulfilled the inclusion criteria of a documented PHI or <2 years between the last negative and first positive HIV-1 test. Despite this limitation, the trajectories of the biomarkers in our training dataset agreed with previously published estimates obtained using larger datasets ^7,14,23–25^ and this gives us confidence that model design and parameter values are valid for other patients.

The strengths of our approach are that it relies on routine or easy to collect data and provides estimates of the size of undiagnosed population, stratified by HIV transmission group, explicitly accounting for exogenous infections. In addition, the method considers the several sources of uncertainty involved, such as differences in HIV-1 testing behavior, differences in disease progression, and biomarker measurement error. Because the method estimates TI for each investigated person it allows for detailed HIV-1 epidemiological investigations, such as the causes and consequences of late presentation of HIV infection ^30^ and the effectiveness of HIV-1 prevention (e.g. preexposure prophylaxis) ^31,32^. The method can easily be adapted to other biomarkers like the LAg avidity assay and viral diversity ^33–35^.

Our study suggests a way forward for generating up-to-date reports on the key parameters (incidence and number of undiagnosed persons) needed to understand the effectiveness of HIV control methods, the application of public health triage, and the progress towards the UNAIDS 90-90-90 targets. The global movement of people in response to a changing world means that health systems will be challenged with increasing numbers of foreign-born persons; surveillance methods will need to correctly account for infections in those populations that occur both before and after immigration. Integrating computational modules into public health surveillance streams that can account for complex, heterogenous data with missing values will greatly increase the utility of surveillance not only as a passive monitoring tool but as an active intervention aid.

